# Patterned illumination enables denser deep-learning based single-molecule localization microscopy

**DOI:** 10.1101/2025.03.09.642203

**Authors:** Jelmer Cnossen, Shih-Te Hung, Daniel Fan, Josiah B. Passmore, Lukas C. Kapitein, Nynke Dekker, Carlas S. Smith

**Author notes:** these authors contributed equally to this work (Jelmer Cnossen, Shih-Te Hung, Daniel Fan).

## Abstract

Single-molecule localization microscopy (SMLM) improves resolution beyond diffraction limits by imaging cellular structures at the nanometer scale. Recent advances include using modulation patterns to improve localization precision, and deep learning to accurately process high-density samples with overlapping fluorophore emissions, thus improving imaging speed. A method combining these two advances, SIMCODE, is presented here, allowing high-density modulation-enhanced SMLM. SIMCODE achieved resolution improvements in high-density areas compared to SMLM, deep learning-based SMLM (DECODE), and modulation-enhanced SMLM alone (SIMFLUX). In DNA-PAINT imaging of COS-7 cells, SIMCODE showed improvements in the Fourier Ring Correlation and resolution-scaled Pearson coefficient, with overall improvement increasing as imaging buffer concentration increased five-fold. Modulation-enhanced localization microscopy combined with deep learning thus produced higher quality reconstructions at higher emitter densities (i.e., ∼3× the number of detected spots). This will enable faster imaging, higher labeling densities, and more flexibility in fluorophore choice, which are important for studying dynamic processes and densely labeled structures.

## Main

Super-resolution fluorescence microscopy has become an indispensable tool in empowering scientists to visualize biological systems at subcellular and even single-molecule levels. One particularly notable technique in this field is single-molecule Localization Microscopy (SMLM)^1–3^, which facilitates imaging at the nanometer scale. This method involves capturing numerous camera frames in which only a small number of emitters stochastically emit light, preventing their diffraction patterns, known as the point-spread-function (PSF), from overlapping. This approach enables the precise localization of emitter coordinates by fitting a model of the PSF and subsequently reconstructing a super-resolution image from the acquired coordinates. However, this approach depends on having a sparse emitter density and a high photon budget, limiting the temporal resolution when imaging dynamical processes. This results in prolonged acquisition times, leading to low throughput and significant constraints on the stability of the imaging system^4^, as well as low labeling densities and reduced choice of fluorophores.

Recent advances in SMLM have focused on improving spatial precision through the use of modulated illumination patterns, or improving measurement throughput through the use of advanced computational methods to accurately process high density samples with many overlapping fluorophore emissions. In the first instance, the use of modulation patterns can significantly improve localization precision (SIMFLUX^5^, ModLoc^6^, ROSE^7^), with these methods collectively referred to as modulation-enhanced SMLM^8^. Secondly, the steady improvement of fitting algorithms that are able to deal with higher density measurements, where the PSFs of nearby fluorophores start to overlap with each other, has significantly shortened measurement duration. As the reconstruction of the super-resolved image involves both estimating the number of molecules and the position of these molecules, frames with higher densities of fluorophores pose a difficult statistical estimation problem. Methods such as 3B^9^ and BAMF^10^ solved this accurately using Monte Carlo approaches, but the inference itself can require days of computation making it infeasible as practical tools for biologists. While several groups have proposed faster computational methods^11–13^ to solve the overlapping-emitter localization problem assuming 2D localization, key breakthroughs came with the application of deep learning to SMLM: both DECODE^14^, DeepSTORM^15^, and others^16,17^ showed that deep learning approaches are ideally suited to solve these complex inverse problems, performing high density 2D and 3D localization in a practical timeframe.

Nevertheless, standard hardware and analysis algorithms for modulation-enhanced SMLM requires emitters to be both sparse and slow-blinking, limiting measurement throughput and restricting imaging stability beyond even non-modulation-enhanced SMLM. Similarly, deep learning based methods to directly reconstruct emitter numbers and locations have reached performance limits. However, a solution that leverages both approaches simultaneously, combining the spatio-temporal benefits of both methods, remains unexplored. In this work, the combination of modulation-enhanced SMLM with deep learning was studied, and its performance against standard SMLM alone, modulation-enhanced SMLM alone (using SIMFLUX), and SMLM with deep learning alone (using DECODE) was benchmarked. This combination, henceforth described as SIMCODE, simultaneously addresses the challenges of limited imaging speed, labeling density, and localization precision in a systematic manner, with significant performance improvements over SMLM alone, SIMFLUX alone, and DECODE alone.

The optical system used six modulation patterns (two orthogonal orientations and three phases) to complete an imaging cycle. Utilizing a digital micromirror device (DMD) as a diffraction grating, multiple diffraction orders were formed, of which only the +/-1 diffraction orders were transmitted via a spatial filter. Different orientations were achieved by applying distinct DMD patterns, while phase shifts were realized by shifting the DMD pattern. The DMD’s microsecond-level pattern orientation and phase adjustment was well suited for fast switching between six patterns within the limited on-time of a single-molecule in SMLM experiments. A quarter-wave plate adjusted the DMD reflected light to circular polarization, after which a pizza polarizer ensured x- and y-linear polarization for the horizontal and vertical oriented beams respectively. This equalized the intensity between the x- and y-patterns thus maximizing modulation contrast for each orientation. Fine adjustment of the quarter-wave plate allowed fine-tuning of the modulation contrast since the downstream dichroic mirror can subtley modify the polarization state of the beam. This system simplified polarization control by utilizing a passive element (pizza polarizer) instead of an active element (Pockels cell), and achieved a modulation depth exceeding 90% in SMLM imaging (based on the low density experimental dataset). The DMD-based hardware design allowed for fast pattern switching and offered a more streamlined and reproducible system compared to previous work^5^, where modulation patterns were generated with relatively slow piezo stepping and a Pockels cell. Importantly, by using robust optical components to mitigate thermal and vibrational fluctuations, a fan-less DMD to further reduce vibrations, and a short optical excitation path length to mitigate fluctuations in the modulation pattern, this optical system was able to achieve stable imaging with fast, customizable pattern switching (also see Supplementary Fig. S6).

To process high-density raw data from the setup, a deep-learning based method was used, with the aim to achieve high accuracy across a wide range of emitter densities and brightness levels (see Methods). The network design builds upon the pioneering work from DECODE^14^, and was trained to output a Gaussian Mixture Model that describes the distribution of possible emitters and their positions 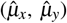, background photon counts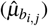, and emitter photon counts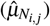. After the raw images were processed by the deep learning model and converted into localizations, the modulation patterns were calculated using the emitter photon counts also estimated by the deep learning model (Fig. 1). The pitch and orientations of the patterns were found using Fourier domain peak finding based on the localization reconstruction^5^. Pattern phases were subsequently retrieved by fitting the sinusoidal modulation pattern to the deep learning estimated emitter photon counts, binned over time to account for phase drift during the measurement (see Supplementary Information). After pattern estimation, the modulation-enhanced positions were estimated by performing a Levenberg-Marquardt^18,19^ based maximum-likelihood fit, fitting each spot’s 6 intensities to the known modulation patterns. The final position estimates were obtained by merging the modulation pattern based position estimates with the deep learning model based position estimates, taking into account their predicted uncertainties. This presented method, SIMCODE, is schematically shown in Fig. 1. For benchmarking, the data was also processed using the SIMFLUX pipeline (i.e., without deep learning) and using DECODE on summed frames disregarding intensity (i.e., without pattern information).

**Figure 1.**
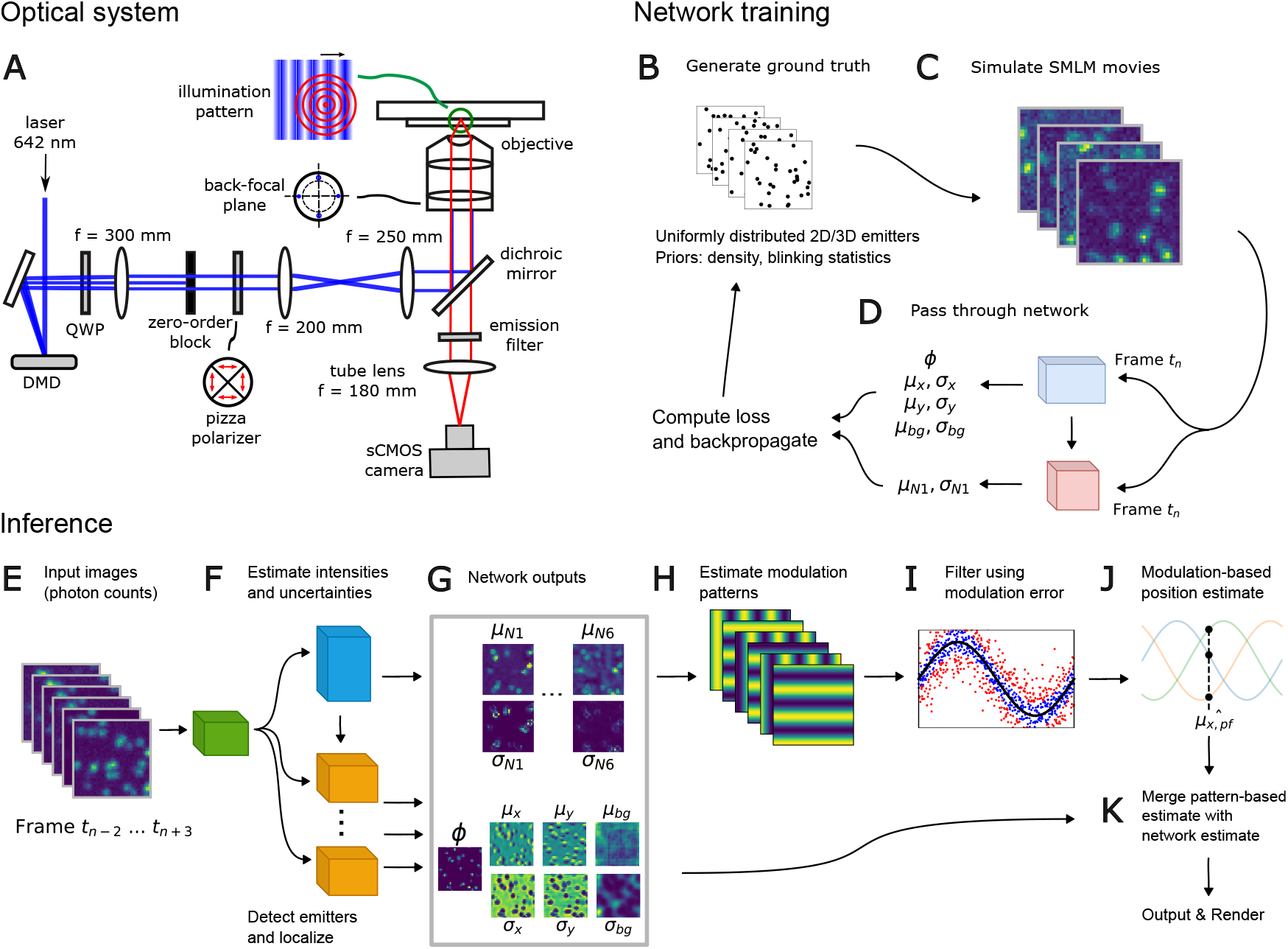
An overview of the SIMCODE processing pipeline. (**A**) The optical system uses a digital mirror device to produce controllable sinusoidal modulation patterns. The deep learning model was trained on simulated SMLM recordings, in which a set of blinking emitters was generated (**B**) and rendered into sets of 24 frames that were combined into larger batches (**C**). Priors for this simulation include the width of the 2D Gaussian PSF, the average density of emitters, the average on-time of emitter (exponentially distributed), and the emitter intensity distribution. The network outputs (**D**), for each pixel, a probability of having an emitter (*ϕ*), and the properties and uncertainties of that emitter (*µ* and *σ*, respectively). See the Methods section for a description of the loss function based on these properties. In inference mode, raw input images (**E**) were converted into localizations and their positions and intensities (**F, G**). The localization intensities were then used for modulation pattern estimation (**H**), and subsequently filtered using the modulation error^5^ (**I**). This removes localizations that were not in their on-state for the full 6 frames. The modulation-enhanced position (**J**) was then estimated by assuming that the estimated intensities have normally distributed errors, and applying a Levenberg-Marquardt routine to find the maximum-likelihood position of the emitter. Finally, the position estimate from the deep learning network (*µ*_*x*_, *µ*_*y*_) was merged with the maximum-likelihood modulation-based estimate (**K**), weighted by their Fisher information, to further improve localization precision.

First, to evaluate the performance of SIMCODE in-silico, simulations of SMLM microtubule measurements were performed. The resulting simulated movies were processed using a regular single-emitter SMLM pipeline, as well as SIMFLUX, DECODE, and SIMCODE (see Fig. 2). Supplementary Fig. S1 and S2 shows an overview of reconstructions of the same ground truth structure with different densities of active emitters. The randomly generated microtubules have some sparse and some denser areas, and the reconstructions shown (Fig. 2A, B, C, E, and F) focus on the densest area, as there the largest differences between methods can be observed. The reconstructed images revealed two prominent artifacts of localization: false positives, where closely situated emitters were erroneously localized as a single emitter resulting in additional localizations between the true emitter positions, and false negatives, occurring in high-density areas where the method failed to detect individual emitters, leading to missing sections of the ground truth structure. Examination of the reconstructions demonstrated that SMLM produced numerous false positives in high-density regions, exacerbated by increasing emitter densities. SIMFLUX exhibited the ability to reject false positive localizations by leveraging modulation pattern information but yielded a sparse reconstruction with a lower number of localizations (Fig. 2D) and thus high number of false negatives. This sparsity was likely due to the requirement for emitters to be isolated for a full 6 frames, impacting intensity and position estimates. Consequently, large portions of the ground truth structure were absent in the reconstruction. Notably, both deep learning methods, DECODE and SIMCODE, mitigated these reconstruction errors while SIMCODE delivered an overall higher Fourier Ring Correlation (FRC, Fig. 2G, 20.7 nm for SIMCODE compared to the next best 28.2 nm for DECODE, for emitter density of 1 emitter per *µm*^2^) across multiple emitter densities (0.5 to 4 localizations per frame per *µm*^2^, Supplementary Fig. S1 and S2) as well as lower total number of false positives and negatives as per the Jaccard similarity index (i.e., 0.89 for SIMCODE compared to 0.68 for DECODE, Fig. 2H).

**Figure 2.**
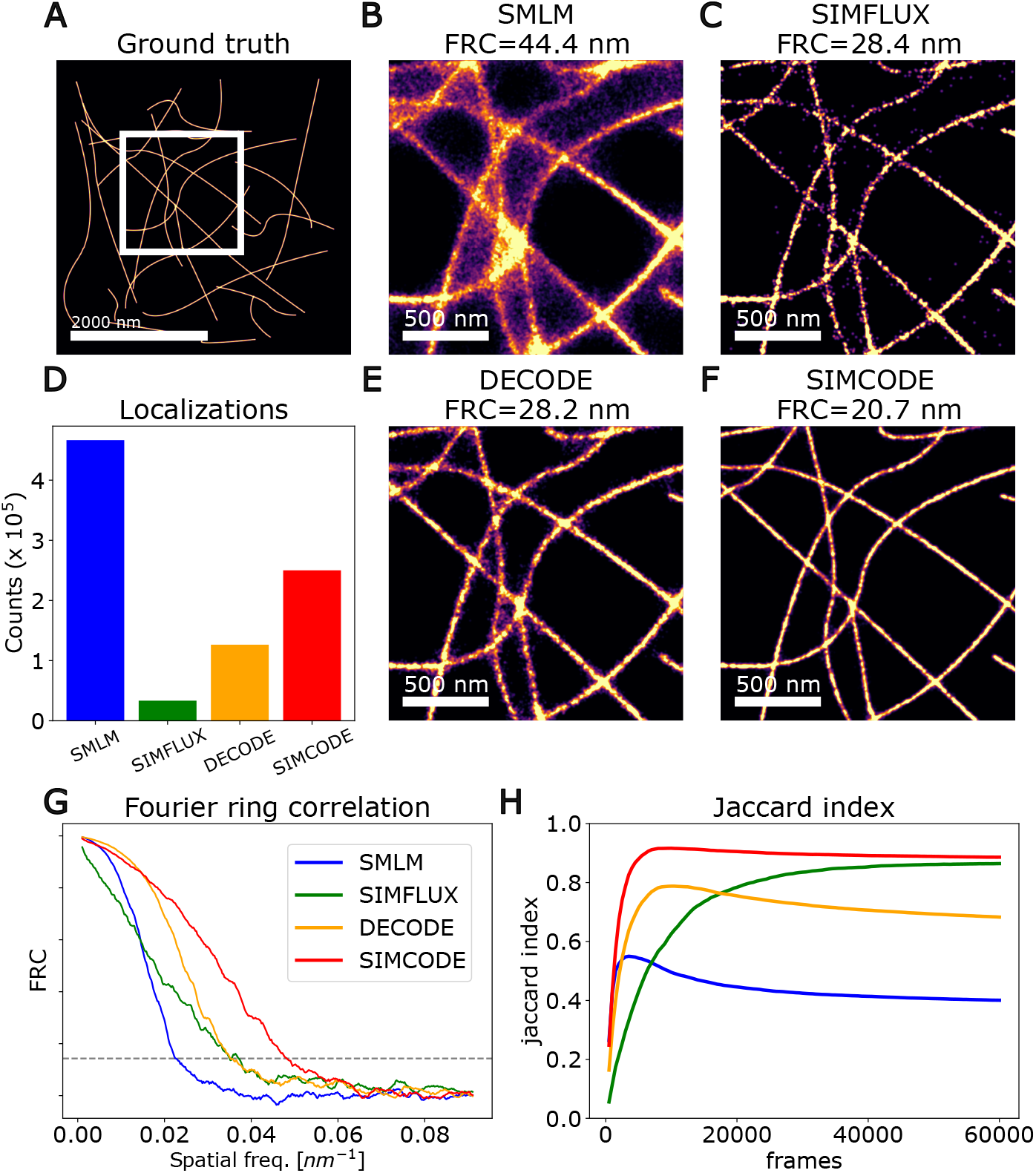
A comparison of several localization methods on simulated dataset with randomly generated microtubules, with modulation patterns applied. (**A**) Ground truth positions of emitters, with blinking probabilities set to result in an average emitter density of 1 emitter per *µm*^2^ per frame. For generation details see Supplementary Information. (**B, C, E, F**) Reconstruction of the zoomed in area marked by the rectangle in (**A**) using (**B**) SMLM, (**C**) SIMFLUX, (**E**) DECODE, and (**F**) SIMCODE. (**D**) Number of localization assignments for each of the different methods. (**G**) Fourier Ring Correlation (FRC) curves for the various methods. The FRC curves were smoothed with a 10-sample uniform filter to obtain more accurate FRC values. (**H**) Jaccard Index for the different reconstruction methods vs. the number of frames.

To analyze the speed at which different methods reconstructed ground truth structures, a simulation of a localization microscopy experiment employing four distinct emitter densities was performed (Supplementary Fig. S2). Each density variation underwent processing using four different reconstruction methods: regular SMLM, SIMFLUX, DECODE, and SIMCODE. The Jaccard similarity index over time was computed for different densities and the four different reconstruction methods. Deep learning methods DECODE and SIMCODE achieved final Jaccard indices significantly higher than SMLM at high emitter density of 4 localizations per frame per *µm*^2^ (0.76 for SIMCODE and 0.60 for DECODE compared with 0.37 for SMLM, and 0.41 for SIMFLUX, see Supplementary Fig. S2). The slight drop in Jaccard index (Fig. 2H) after reaching peak values was attributed to the generation of false positives despite correct ground truth structure identification by the reconstruction. Importantly, SIMCODE outperformed DECODE overall in terms of Jaccard index (Fig. 2H and Supplementary Fig. S2) and was able to reach its maximum performance (i.e., its maximum Jaccard index) faster at higher emitter densities (Supplementary Fig. S2). This showed that SIMCODE can deliver improved spatial resolution compared to SMLM, SIMFLUX, or DECODE more quickly (i.e., with no significant change in Jaccard index of ∼0.8 at 10× the number of localizations per frame, see Supplementary Fig. S2). Finally, note that although SIMFLUX showed a steady increase in Jaccard index with the number of frames being imaged, it needed many more frames and thus more time to achieve a similar level of reconstruction.

To assess the performance of SIMCODE on experimental data, the performance of the new optical design was first characterized, which showed improved modulation contrast (>90%) and pattern speed (purely camera and exposure time limited). Second, African green monkey kidney COS-7 cells were imaged using the optical system for two different concentrations of labeling dyes. Fig. 3 (0.5 nM image buffer concentration) and Fig. 4 (2.5 nM image buffer concentration) show the reconstructions on experimental data for SMLM, SIMFLUX, DECODE, and SIMCODE, at two different emitter densities. The average number of spots detected per imaging cycle for 0.5 nM and 2.5 nM image buffer concentration specimens were 63.7 and 190 respectively, an increase of 3× (see Fig.3K and Fig.4K). SIMCODE was shown to deliver improvement over SIMFLUX and DECODE through the fusion of modulation-enhanced localization microscopy with high-density deep learning-based SMLM. Especially in high-density regions, an enhanced localization density and localization precision for SIMCODE compared to SIMFLUX and DECODE was observed. At low image buffer concentrations, SIMCODE showed an 18% FRC improvement over the next best method (SIMFLUX) at triple the number of localizations. SIMCODE also showed a 29% FRC improvement over DECODE with a similar number of localizations (Fig. 3J, K) as well as a higher overall resolution-scaled Pearson (RSP) correlation coefficient^20^ (i.e., 0.64 for SIMCODE compared to the next best DECODE with 0.61, Fig. 3L). Importantly, at high image buffer concentrations, the FRC for SIMCODE was 15% better than the next best DECODE with additionally more than double the number of localizations than DECODE as well as almost double the RSP coefficient compared to DECODE (Fig. 4J, K, and L). Concurrently, at high image buffer concentrations, the RSP coefficient for SIMCODE remained high, indicating that image quality was preserved even at high emitter density. On the other hand, the RSP coefficient for both SMLM, SIMFLUX, and DECODE, decreased at high emitter densities (i.e., SMLM from 0.4 to 0.09, SIMFLUX from 0.54 to 0.12, and DECODE from 0.61 to 0.46, while SIMCODE increased from 0.64 to 0.75, as image buffer concentration increased from 0.5 nM to 2.5 nM respectively, Fig. 3L vs. Fig. 4L). Meanwhile, the number of localizations for SIMFLUX and DECODE decreased as image buffer concentration increased (∼3×10^5^ to ∼1×10^5^ and ∼1×10^6^ to ∼7×10^5^ respectively, Fig. 3K and Fig. 4K), which, combined with the decrease in RSP coefficient, can indicate an increased number of false negatives. Similarly, the number of localizations for SMLM increased as image buffer concentration increased (∼2.8×10^6^ to ∼3.9×10^6^), which, combined with the decrease in RSP coefficient, can indicate an increased number of false positives. On the other hand, the number of localizations for SIMCODE increased (∼9×10^5^ to ∼1.7×10^6^) together with an increased RSP coefficient as image buffer concentration increased, which can indicate an increased number of accurate localizations. SIMCODE thus showed overall better image quality at high emitter density, where other methods exhibited deteriorating performance.

**Figure 3.**
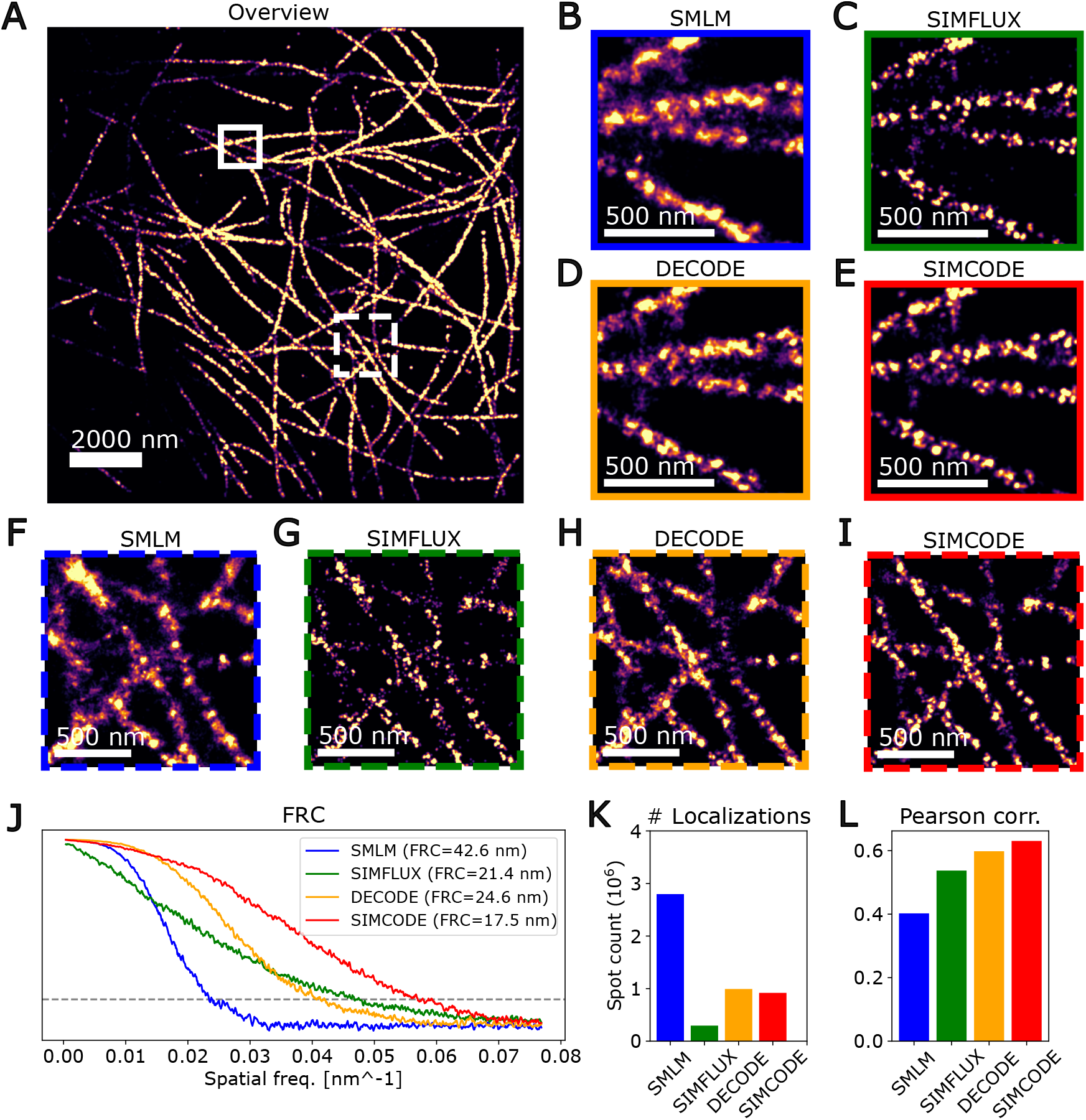
DNA-PAINT imaging of tubulin in African monkey kidney COS-7 cells (see Methods) at a 0.5 nM imager concentration, which causes a relatively low number of active emitters in the field of view (average of 63.7 spots per imaging cycle). Reconstructions are done using the 4 different methods as described in the Methods section. The overview in (**A**) shows SIMCODE reconstructions, whereas the other methods are visualized in zoom-in panels **B-E** and **F-I**. All localizations with an uncertainty (either Cramer-Rao Lower Bound or *σ* from Gaussian Mixture Model) lower than 0.15 pixels were filtered out for all methods, including FRC calculation. (**J**) FRC curves, calculated on the full reconstructions. (**K**) Localization counts, calculated after filtering. (**L**) Resolution-scaled Pearson (RSP) correlation coefficients.

**Figure 4.**
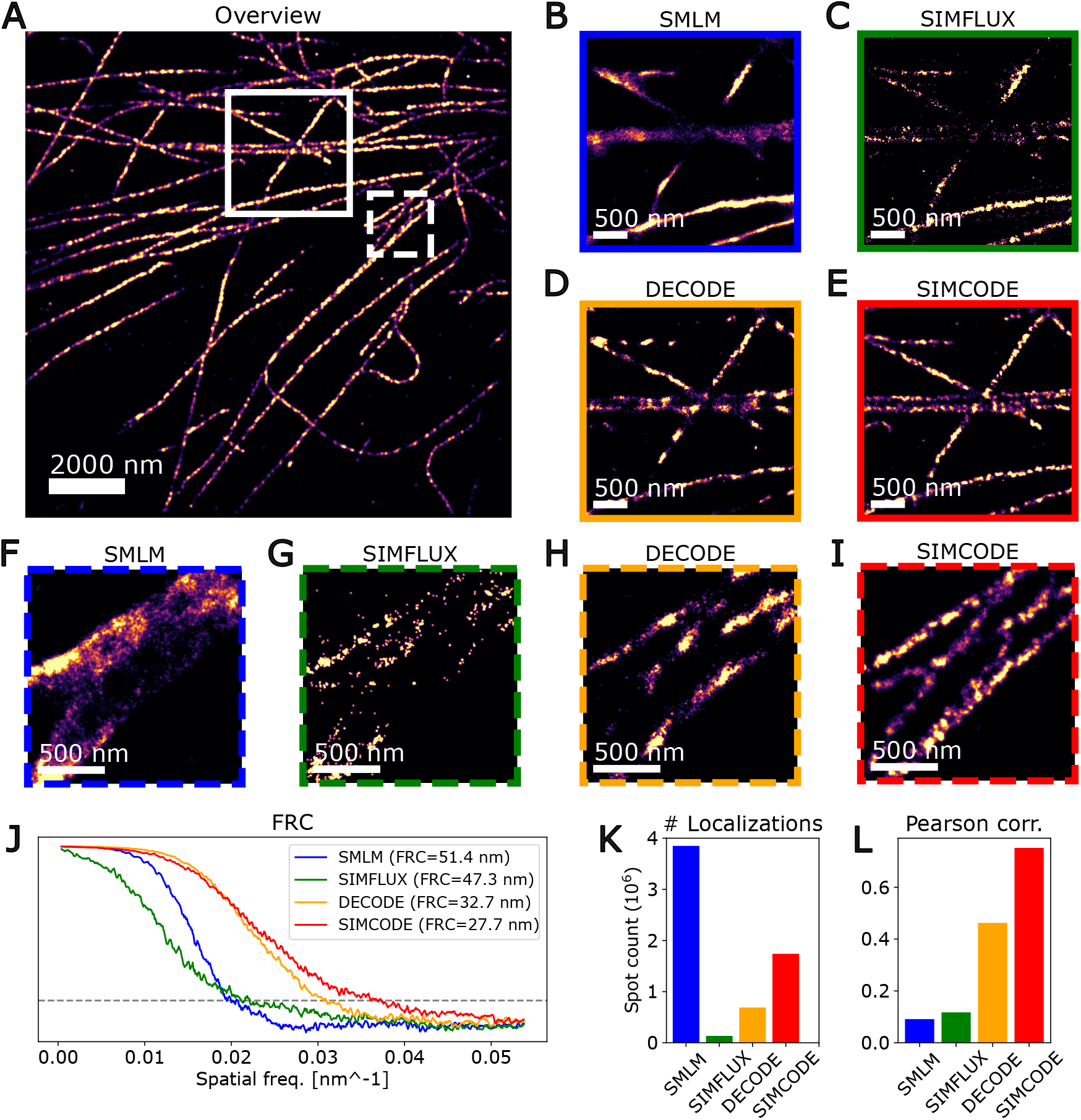
Reconstructions of an experimental recording of DNA-PAINT imaging of African monkey kidney COS-7 cells using a 2.5 nM imaging buffer, resulting in a higher density of emitters relative to Fig. 3. An average of 190 spots per imaging cycle were found, as detected by the deep learning model before any pattern-based filtering. Data processing and recording was identical to Fig. 3.

In conclusion, a super-resolution microscopy modality combining modulation-enhanced SMLM with deep-learning based SMLM (i.e., SIMCODE) was successfully demonstrated. Both simulations and experimental results show that SIMCODE was able to maintain the precision benefits of SIMFLUX while imaging at a much higher emitter density where traditional SMLM and SIMFLUX produce a significant number of false positives and false negatives. Simulations across various emitter densities and experimental results demonstrated the spatio-temporal improvements of SIMCODE, offering enhanced localization density and precision compared to existing methods. Results show that, in addition to improving the localization precision of individual emitters, modulated illumination also provides an effective way of filtering out false positives, in turn allowing an increased labeling density. Increased labeling density enables faster imaging, higher fluorophore density, and greater flexibility, which are beneficial for studying dynamic processes and densely labeled structures (see Supplementary Fig. S8). SIMCODE offers benefits similar to DECODE, with added advantages of higher throughput, reduced reliance on instrument stability, and less stringent fluorophore preparation requirements. Unlike other deep-learning-based SMLM approaches, SIMCODE integrates deep learning-derived emitter intensity and position information with a known modulated illumination pattern. This combination enhances imaging performance, particularly in high-density areas, surpassing the capabilities of traditional SMLM, deep-learning SMLM, or modulation-enhanced SMLM used independently. These advancements address key limitations in traditional SMLM, such as imaging speed and labeling density. By leveraging modulation-enhanced techniques combined with deep learning methodologies, SIMCODE represents a crucial step towards overcoming challenges associated with imaging dynamic biological systems at the sub-cellular and molecular levels.

## Methods

### Experimental setup

The laser source in the experimental system was a 642 nm continuous-wave, single-mode laser (2RU-VFL-P-2000-642-B1R, MPB, Pointe-Claire, Canada) with 2 W maximum output power. The intensity of the laser was controlled by an acousto-optic tunable filter (AOTF, AOTFnC-400.650, AA OPTO-ELECTRONIC, Orsay, France). The laser was then coupled into a polarization-maintaining single mode fiber (PMSF, Thorlabs, Newton, U.S.A.) using an objective lens (UMPlanFLN 20X NA 0.5, Olympus, Tokyo, Japan). A fiber port (SM1SMA, Thorlabs), which holds the PMSF, was mounted on a translation lens mount, by which fiber coupling efficiency could be optimized. To match the polarization direction of the PMSF and the laser, a half-wave plate (WPH10M-633, Thorlabs) was placed between the AOTF and the PMSF to rotate the polarization direction of the laser. The half-wave plate was mounted on a rotation mount (RSP1×15/M, Thorlabs), which offers a degree of freedom to change the fast axis angle of the half-wave plate. The laser was then coupled out from the PMSF via an objective lens (PlanN 4× NA 0.1, Olympus). Then, the beam passed through a high extinction ratio Glan-Taylor calcite polarizer (GT15-B, Thorlabs) to ensure the purity of the polarization state of the laser. A 2× demagnification relay system consisting of two achromatic lenses (f = 200 mm and f = 100 mm, AC254-200-A and AC254-100-A respectively, Thorlabs) was then adopted to reduce the laser spot size. Afterward, a protected silver Zerodur mirror (PF1011-P01, Thorlabs) was used to reflect the laser to a digital micro-mirror device (DMD, Vialux, V-7000 VIS, Chemnitz, Germany). The DMD works as a programmable amplitude optical grating, by which the grating pattern could be rapidly switched to realize patterns for modulation enhanced imaging. The maximum switching rate of the DMD pattern was 22 kHz based on vendor specifications. If the blinking time of a single-molecule is assumed to be roughly 50 ms, the system will need to switch between 6 patterns within a blinking time of 50 ms, which is equivalent to a 120 Hz DMD switching rate. Therefore, the switching rate of this DMD was fast enough to realize the imaging presented in this work. Then, another protected silver Zerodur mirror was used to align the DMD output beam towards the excitation path. A quarter-wave plate was placed immediately after this mirror to ensure a circularly polarized beam such that the intensities of x- and y-modulation patterns were equal. Then, an achromatic lens (f = 300 mm, ACT508-300-A, Thorlabs) was used to transform the diffracted light into the Fourier plane. A pinhole was placed in the Fourier plane to block the zero-order diffraction light, followed by a pizza polarizer (VIS 600 BC5, Codixx, Barleben, Germany) to ensure the purity of desired s-polarization state at the back-focal plane of the objective lens for both x- and y-oriented patterns. The extinction ratio of the pizza polarizer was ∼10^6^ at 640 nm. Solely s-polarized light will result in a high modulation contrast^5^ at the sample. Then, a 4f system consisting of two achromatic lenses (f = 200 mm and f = 250 mm, ACT508-200-A and AC254-250-A respectively, Thorlabs) was used to relay the Fourier plane to the back-focal plane of the imaging objective lens. A 3-mm-thick dichroic mirror (R405/488/532/635 *λ* /5 flat, Semrock, Rochester, U.S.A.) was used to separate the excitation and emission light. The thickness ensures lower aberrations from the reflection surface of the dichroic mirror, avoiding the problem that the optimal modulated illumination pattern in x- and y-directions appear at different axial positions and minimizes the light distortion of the modulation patterns. A high numerical aperture (NA) total internal reflection fluorescence (TIRF) objective lens was used as an imaging objective lens (UPLAPO60XOHR, NA = 1.49, Olympus), which allows for TIRF structured illumination microscopy (TIRF-SIM) modality. The DMD was used as a periodic binary grating with an effective pitch size of 54.5 *µ*m, resulting in an illumination NA of 1.47 at the back-focal plane of the imaging objective lens. In the emission path, an emission filter (FF01-446/510/581/703-25, Semrock) was used to block the excitation laser. A tube lens (TTL180-A, Thorlabs) was then used together with the imaging objective lens to form a 60× magnification microscope. A sCMOS camera (Zyla 4.2, Andor, Belfast, U.K.) was used to acquire the images. To synchronize the device, an Arduino microcontroller (Uno Rev3, Arduino, Monza, Italy) was used as a master device to trigger the DMD patterns and sCMOS camera image acquisitions.

### Sample preparation

In the DNA-PAINT cell sample, African green monkey kidney fibroblast-like COS-7 cells (CRL-1651, ATCC, Manassas, U.S.A.) were cultured in DMEM high glucose medium (DMEM-HPSTA, Capricorn Scientific, Ebsdorfergrund, Germany) supplemented with 9% fetal bovine serum (35-079-CV, Corning, Corning, U.S.A.) and 1% penicillin/streptomycin (GIBCO 15140-122, Thermo Fisher, Waltham, U.S.A.), at 37°C in 5% CO_2_. Cells were plated on coverslips (Ø18 mm, no. 1.5H, 0117580, Marienfeld, Lauda-Koenigshofen, Germany) 24 hours prior to fixation. Cells were extracted using pre-warmed 0.35% Triton X-100 (X100, Sigma-Aldrich, St. Louis, U.S.A.) +0.2% glutaraldehyde (G7526, Sigma-Aldrich) in MRB80 (80 mM K-Pipes, 1 mM EGTA, 4 mM MgCl_2_, pH 6.80) for 1 min, and then fixed using 4% paraformaldehyde in 1×PBS for 10 min at 37°C. Following fixation, cells were washed 3 times in 1×PBS for 15 min, permeabilized using 0.2% Triton X-100 for 10 min, washed again, and blocked using 3% bovine serum albumin (8076.3, Carl Roth, Karlsruhe, Germany) in 1×PBS. The primary antibody (rabbit anti-alpha tubulin, ab52866, 1:1000 dilution, Thermo Fisher) was diluted in blocking buffer and incubated for 2 hours. Following primary incubation, cells were washed again in 1×PBS, and processed for DNA-PAINT using the Massive-AB 3-plex labelling kit (Massive Photonics, Munich, Germany). Cells were washed twice in 1×washing buffer for 5 min, then incubated with the secondary antibody (Anti-Rabbit IgG + Docking site 2, 1:500 dilution) diluted in antibody incubation buffer for 1 hour, before washing again three times in 1×washing buffer for 15 min. The imaging buffer of the DNA-PAINT sample consisted of the CF660 DNA-PAINT imager and the dilution imaging buffer. By varying the imaging buffer concentration, the emitter density could be adjusted for single-molecule imaging. A 0.5 nM imager concentration was used for a low emitter density dataset, which was processed using previously published SIMFLUX code^5^, and a 2.5 nM imager concentration was used for a high emitter density dataset, which could only be processed by the proposed SIMCODE method in this work. The imaging buffer was applied onto a glass slide with a single cavity (Thorlabs, MS15C1), and the coverslip was then affixed to the glass slide with the cell side facing downwards, ensuring the cell was fully immersed in the imaging buffer. A two-component gel (Twinsil, Picodent, Wipperfürth, Germany) was used to seal the coverslip and avoid oxygen exchange.

### Data acquisition

For data acquisition, Andor SOLIS software was to control the camera. The image readout was in 16-bit mode and the camera was under the control of an external triggering signal. The DMD pattern updates 6 times (2 axes and 3 phase steps) to form a complete imaging cycle. 20000 imaging cycles were recorded for the low density dataset, and 30000 imaging cycles for the high density dataset (120k and 180k frames, respectively). The imaging window was 120×120 pixels and the exposure time was 5 ms. The total exposure time of an imaging cycle was 30 ms.

### Method comparison

The SMLM reconstructions were done using a conventional pipeline of spot detection, region-of-interest (ROI) extraction, and a Levenberg-Marquardt maximum-likelihood fit of isolated emitter images. A ROI size of 8×8 pixels was used for this (for a 2D Gaussian PSF of *σ* = 1.3 pixels). A *χ*^2^ filter was applied to the fitting results, by comparing the expected value of pixels in the emitter ROIs with the true pixel values, to attempt to remove bad fits and overlapping emitters. As the SIMFLUX and SIMCODE simulations and experimental data consisted of modulated frames, these were summed over a moving window of size 6, before passing them to SMLM and DECODE. While this ensures that the two methods were given emitter images with equal photon counts, summing them will also effectively increase emitter density for these two methods. However for the alternative, i.e., feeding modulated images into the SMLM and DECODE pipeline, a lower image quality was found, so the summed window approach was used to compare methods. The DECODE pipeline was trained to use a 3-frame context window, so in terms of input pixel data, the DECODE network has access to 18 modulated frames. DECODE was modified to swap the X and Y coordinates to match the rest of the code. In additional, a 2D Gaussian PSF was used during model training, instead of a 3D cubic spline, so the PSF model was exactly the same for SMLM, SIMFLUX, DECODE, and SIMCODE processing. For the experimental datasets, drift correction was done using Drift Estimation by Minimum Entropy^21^ and further detailed in Supplementary Fig. S4 and S7 In order to focus the comparison on the localization method, the drift traces were estimated using the SIMCODE localizations, and applied to the localizations of all methods.

### SIMCODE architecture

The network architecture of SIMCODE is illustrated in Supplementary Fig. S3. The deep learning model was largely a modified version of the DECODE architecture, and consists of 3 U-Net^22^ modules and several separate convolutional layers. The first U-Net runs a first pass on the raw frame data, converting it to a latent space. The second U-Net does detection and estimation on a moving window of 6 frames, and the third U-Net estimates spot intensities based on a latent space representation of the detected emitters and their positions. The loss function was similar to DECODE’s, and designed to make the network output a Gaussian mixture model (GMM) that maximizes the likelihood of sampling the ground-truth set of emitters. In short, for each pixel of each frame, the model outputs estimated parameters *θ* = (*x, y, N*_1,1_, …, *N*_2,3_, *b*_1,1_, …, *b*_2,3_) and the estimated uncertainties, where *x, y* is the 2D position, *b*_*i, j*_ is the background (in photons/pixel), and *N*_*i, j*_ the emitter intensity (in number of photons) at the *i*-th angle and the *j*-th phase step. This was implemented using a GMM resulting in a mean *µ*_*θ*_ and covariance *σ*_*θ*_, which was assumed to be diagonal. The model was trained to predict emitters at each frame, using the pixel data from a moving window around the current frame (the preceding 3 frames, the current frame, and the proceeding 2 frames). Key differences with DECODE’s loss function are the introduction of multiple intensities per emitter, and that the background fluorescence was also included in the GMM parameters and not estimated with a separate least squares loss. The model configuration used here is by no means optimized for compute speed or memory use. The layer channel sizes are chosen high enough to not create a bottleneck for the localization performance, in order to test the efficacy of deep learning in modulation enhanced SMLM. For a complete definition of the loss function, see Supplementary Information section 1.1.

### SIMCODE training

The initial step involved training the deep learning model for localization using a known PSF model and rough estimates of the ranges of emitter on-time, intensities, and background fluorescence values. The network was trained on simulated recordings of SMLM experiments, in which fluorophores have uniformly distributed random positions. This ensured that the network was not biased towards any particular structure, and only has the emitter density, PSF model, fluorescence brightness, the range of background fluorescence values and blinking statistics as priors. Training was conducted on a server with Intel i9-10900X and nVidia RTX 3090 24GB running Ubuntu, CUDA 11.8, Python 3.11, and PyTorch 2.2.1. The batch size was optimized to fill GPU memory, but training could be adapted to less powerful GPUs by adjusting batch size and learning rates. Training, performed over 200 epochs. Training employed large sets of small images (32×32 pixels) due to the quadratic scaling of the loss function in compute complexity. However, the resulting network, thanks to the use of convolutional layers, could be applied for inference over arbitrary-sized images. See Supplementary Information for more details on training data generation.

### Data processing

The data processing pipeline is depicted in Fig. 1. The raw images were first converted to photon counts by a fixed gain and offset correction, and then processed by the deep learning model using a moving window of 6 frames. Due to the estimation of spot intensities per frame and per window, the SIMCODE deep learning model was more computationally intensive than DECODE. The 3090 RTX graphics card used processed the frames from Figure 3 in 23.19 min vs. 3:07 min, for SIMCODE and DECODE respectively. Once emitter intensities and variances 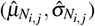 were obtained for each angle (*i* ∈{1, 2}) and phase step (*j* ∈ {1, 2, 3}) from the deep learning model, they were utilized to infer a position estimate. This inference was achieved by performing a maximum likelihood fit using Levenberg-Marquardt, with the estimated intensities and a known modulation pattern (*e*_*i, j*_(*θ*)):

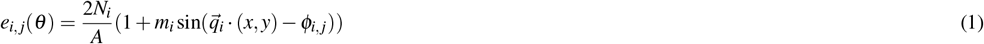

where *ϕ*_*i, j*_ is the modulation phase for angle *i* and phase *j*,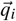 is the modulation frequency for angle *i, N*_*i*_ is the emitter intensity for angle *i, m*_*i*_ is the modulation contrast of the modulated illumination pattern for angle *i, A* is a normalization factor, and *θ* = (*x, y, N*_1_, *N*_2_) with *x, y* as the unknown 2D emitter position. Emitter intensities were independently fitted per modulation pattern angle to account for potential differences in modulation intensity over the field of view, depending on the modulation angle. The estimated photon count as produced by the deep learning model 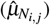 was assumed to follow a normal distribution^23^, where the standard deviation 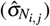 was also estimated by the deep learning network. This leads to the following optimization problem for finding the modulation-enhanced position:

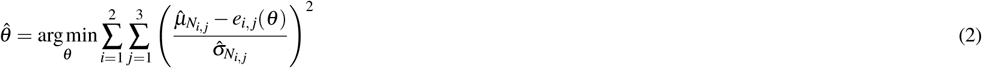

Levenberg-Marquardt (LM)^18,19^ was used for the maximum likelihood estimation, as it does not require computing the second derivative and is potentially more stable than alternatives like Newton-Raphson. A version modified for Poisson distributed samples^24^ is commonly applied to camera pixel data in localization microscopy, but the fluorescence intensities here were estimated from many emitter and background photons, and can be well approximated with a normal distribution. For this reason, least-squares based LM was used here. To implement LM, the residual error was taken from Eq. 2. The (*i, j*)-th element of the matrix representing the residual errors is given by:

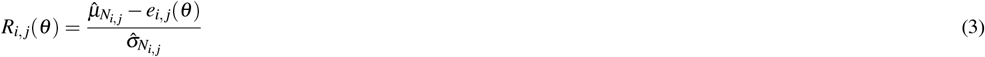

The LM step was then calculated towards the optimum by:

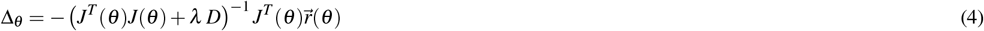

with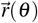 as the vectorization 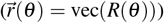 of *R*(*θ*) and *J*(*θ*) as the Jacobian, where 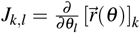such that the derivative of the *k*-th element of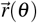to *l*-th element of *θ* was taken. *D* is a damping matrix, with diagonal elements set to diag(*J*^*T*^ (*θ*)*J*(*θ*)), rescaled to have a mean value of 1. *λ* is a scalar damping constant set to 100. From the data, an estimate was obtained of the position without the a priori information of the pattern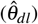. From the intensity estimates, a position estimate was obtained based on the a priori information of the pattern (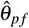, see Supplementary Fig. S5). To combine the position estimate based on both sources of information, the minimum variance unbiased (MVU) estimate of the form:

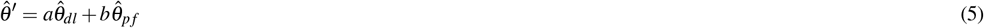

was used, where *a*, and *b* were to be determined. The MVU estimate needs to satisfy two criteria. The first is that the estimate is unbiased: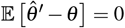. The first condition is easily satisfied because the expected value of the deep learning position estimate is equal to the true position value 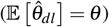 and this is also the case for the position estimate based on the intensities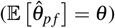. This results in the following expression:

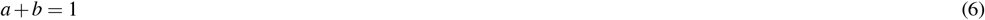

The second is that the estimation variance is a minimum:

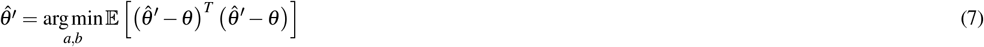

Because the off diagonal components are small, they can be neglected. This simplifies the solution of the MVU estimate to:

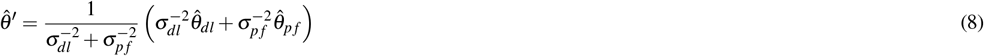

To obtain the MVU estimate, an estimate of the uncertainty is also needed. Typically this can be accurately estimated by the algorithm either through the CRLB^25^ or through a GMM (see Supplementary Information). The deep learning network outputs the uncertainty for the position (*σ*_*dl*_) via the GMM. For the position estimate based on the pattern and intensity estimates the CRLB was used.

### Data analysis

In addition to rendering reconstructions, the Fourier Ring Correlation (FRC)^26^ was assessed for each reconstruction. FRC, a widely used metric for comparing image resolution in localization microscopy, was computed for Fig. 2, 3, 4, and Supplementary Fig. S2. However, as observed in reconstructions, while FRC is valuable for assessing overall image resolution, it does not appear to change meaningfully when large portions of the image are blurred or left out. An alternative to the FRC, and to evaluate how fast the different localization methods can reconstruct the ground truth structure, the Jaccard similarity index was computed between the reconstructions and the ground truth structure:

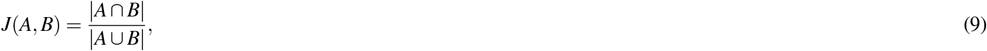

where *A* and *B* are sets containing 2D coordinates, with *A* being the localizations and *B* the points from the ground truth structure. For experimental data where the ground truth was unknown, the resolution-scaled Pearson (RSP) correlation coefficient^20^ was used to indicate the quality of the reconstructed image. The RSP is defined as the Pearson correlation coefficient between the pixels values of the average measured image, versus the pixel values of the super-resolution reconstruction blurred by the known PSF. The result is a coefficient between -1 and 1, where a RSP coefficient towards 1 indicates a better quality reconstruction.

## Acknowledgements

J.C., S.H., D.F. and C.S.S. were supported by the Netherlands Organisation for Scientific Research (NWO), under NWO START-UP project no. 740.018.015 and NWO Vidi project no. 20390. D.F. thanks Dimitri Kromm for assistance with the optical setup. D.F. was partly supported by University of Melbourne’s Early Career Researcher Grants Scheme 2024.

## Data availability

Experimental data files and model weights can be found on https://doi.org/10.5281/zenodo.14642059 Other processed data files are available upon request.

## Code availability

Software for processing the datasets is available as Supplementary Software at https://github.com/jcnossen/ simcode, and includes a Google Colab notebook for an example of processing simulated data, as well as setting up network training.

## Author contributions statement

J.C. and C.S.S. conceived the experiment, J.C wrote the localization pipeline code, J.C., S.H., and D.F. conducted the experiment, J.C. and S.H. analysed the results. All authors reviewed the manuscript.

## Notes

### Competing Interest Statement

The authors have declared no competing interest.

