## supplementary notes for "Patterned illumination enables denser deep-learning based single-molecule localization microscopy"

### 1 Deep learning implementation

#### 1.1 Training and loss function

Training of the network was done on randomly generated emitters with a uniform random distribution over a 32 × 32 pixel grid. The network performance was evaluated after each epoch, including monitoring of the network localization precision on single isolated emitters, to check if it correctly approaches the Cramer-Rao Lower Bound (CRLB).

Spot brightness values were generated from a log-normal distribution with a mean of 600 photons per frame, and a mode of 400 photons per frame. This ensures that most of the training samples have spots close to the real data, but there is a long tail up to 10000 photons to make the network more robust for high density, bright areas.

Modulated patterns were not used in the training phase. Instead, to make sure that the network was able to infer spot intensities for each frame, emitters were generated that have strong frame to frame intensity fluctuations, i.e. $N_f = N(1+\eta_f)$, with fluctuation noise $\eta_f \sim \mathcal{N}(0, 0.5)$, and $N_f$ being the intensity of a spot at frame $f$ and $N$ being the mean spot intensity.

In comparison with DECODE, the SIMCODE loss consists of a Gaussian Mixture Model (GMM), loss component $L_{\text{gmm}}$, and emitter count loss $L_{\text{count}}$. DECODE's background loss is omitted, and instead the background is included in the GMM.

$$L = L_{\text{gmm}} + w_{\text{count}} L_{\text{count}} \tag{1}$$

The probability density of this GMM is expressed as:

$$p_{\text{gmm}}(\theta) = \sum_{i=1}^{W^2} \frac{\phi_i}{\sum_i \phi_i} \prod_{k=1}^{K} \frac{1}{\sigma_{i,k}^2 \sqrt{2\pi}} \exp\left(-\frac{(\mu_{i,k} - \theta_k)^2}{\sqrt{2}\sigma_{i,k}}\right) \tag{2}$$

Here, $\theta$ is a parameter vector with $K$ elements, representing the ground-truth emitter properties: $x, y, N_{1,1}, \ldots, N_{A,B}, b_{1,1}, \ldots, b_{A,B}$, with $A, B$ being the angles and phase steps, respectively. The estimate of each emitter property is treated as an independent normally distributed variable for simplicity and speed. $\phi_i$ is the un-normalized emitter probability at pixel $i$. $W^2$ is the number of pixels in a frame, and $\mu_{i,k}$ and $\sigma_{i,k}^2$ are the network-predicted values for the mean and variance of the emitter properties. In practice, the loss is calculated in log-space due to limited floating-point calculation precision. Given $E$ as the estimated number of emitters, $L_{\text{count}}$ is the emitter count loss that trains the network to output a 1 if a pixel contains an emitter, further described by[1]:

$$L_{\text{count}} = -\log P(E|\mu_{\text{count}}, \sigma_{\text{count}}^2) = \frac{1}{2} \frac{(E - \mu_{\text{count}})^2}{\sigma_{\text{count}}^2} + \log\left(\sqrt{2\pi\sigma_{\text{count}}}\right), \tag{3}$$

where $\mu_{\text{count}} = \sum_{k=1}^{K} p_k$ and $\sigma_{\text{count}}^2 = \sum_{k=1}^{K} p_k(1-p_k)$

The $L_{\text{count}}$ is weighted by $w_{\text{count}}$ (set to 0.01), to improve early training performance. The GMM loss, expressed for a single frame with $E$ emitters, is:

$$L_{\text{gmm}} = -\sum_{j=1}^{E} \log p_{\text{gmm}}(\theta_j) \tag{4}$$

Training was done using the Lion[2] optimizer.

### 2 Simulation of a microtubule dataset

The simulations, the results of which are shown in the main text Fig. 2 and Supplemental Fig. 1 and Fig. 2, were configured to resemble the experimental imaging conditions. A Gaussian point spread function (PSF) model with a standard deviation ($\sigma$) of 1.3 pixels and a pixel size of 108 nm was used to simulate a 40 × 40 pixel area with a total of 60,000 frames.

Filamentous structures resembling microtubules were simulated where ground truth emitter positions were generated by cubic spline interpolation between random points. The ground truth consisted of 23787 generated points, evenly spaced along randomly positioned splines. Binding sites were uniformly distributed along these filaments, with a separation distance of 2 nm. Fluorophores at the ends of these linkers exhibited stochastic blinking, with an average on-time of 6 frames and simulated at 0.1 frame intervals, with emitters having a fixed probability of turning on and off during the intervals. The emitter state transition probabilities $k_{\text{on}}$ and $k_{\text{off}}$ were fixed, with $k_{\text{on}}$ calculated from the desired density and $k_{\text{off}}$ set to $1/6$ such that emitters have an average on-time of 6 frames. The on-state transition probability $k_{\text{on}}$ is given by:

$$k_{\text{on}} = k_{\text{off}} \frac{p_{\text{on}}}{1 - p_{\text{on}}} \tag{5}$$

where $p_{\text{on}} = \rho A/N$, with density $\rho$, area $A$, and $N$ the number of ground truth binding sites.

The illumination pattern was shifted in three discrete phase steps and two orthogonal directions over a pitch of 224 nm, with a modulation depth of 0.9 along both the x- and y-axes to replicate experimental settings. The pixel size was 108 nm. Emitter photon counts were 500 detected signal photons per spot summed over 6 frames and 2.5 background photons per pixel per frame.

#### 3 Modulation based localization

##### 3.1 Pattern estimation

The modulation angle and pitch based on a dataset consisting of localizations with parameters $\theta_x, \theta_y, \theta_{N_i}$, with $i$ being the exposure index was found. Reconstructions were generated at each exposure index for an image at 6× the original resolution, using the modulated intensities $\theta_{N_i}$. Then, the Fast Fourier Transforms of these were computed, the absolute values were taken, and for each modulation angle the 3-phase-step Fourier spectra were summed. Peaks were then detected in these 2D images, and a Discrete Fourier Transform (DFT) was done around the peak, computed directly from the localizations and their modulated intensities. Finally, a quadratic fit was done in both x- and y- directions on the DFT images, to find a precise pitch and modulation angle.

To estimate modulation phase over time, the localizations were divided into blocks of at least 5000 localizations (the last block was larger), and then used to compute the phases and modulation depths over time[3]. This unfortunately underestimated the modulation depth in high density datasets, which is why it was fixed at 0.9 (determined from sparse nanoruler datasets).

Because the digital mirror device (DMD)-based modulation is pixel-based, phase steps should be constant between them, with the phase fluctuations only caused by drift that is shared between all 3 phase steps. This allows a reduction in phase estimation noise by combining the 3 phase step traces into a single phase drift (Supplementary Fig. 4). Thus, after estimating the phase over time for each phase step (as shown in Supplementary Fig. 5), the phase drift for each modulation angle was determined by computing the average of the 3 phase step traces. To validate the estimation, the phase estimation precision was quantified by calculating the standard deviation (Supplementary Fig. 4). Experimental results (see Supplementary Fig. 5) on DNA-PAINT show a typical phase estimation root-mean-square displacement of $< 3$ degrees.

##### 3.2 Modulation based filtering of localizations

To filter out localizations that were not active for a full 6 frames, or otherwise have incorrect or unexpected fluorescence intensities, a modulation error metric was utilized that describes deviation from the expected normalized spot intensity:

$$\gamma = \max_{i,j} \left| \frac{\mu_{N_{i,j}}}{\sum_{i=1}^{2}\sum_{j=1}^{3}\mu_{N_{i,j}}} - e_{i,j}(\theta) \right| \tag{6}$$

Here, $e_{i,j}(\theta)$ models the modulation sine wave at exposure step $i$:

$$e_{i,j}(\theta) = \frac{1}{6}(1 + m_i \sin(\vec{q}_i \cdot (x,y) - \phi_{i,j})) \tag{7}$$

Here, $m_i$ is the modulation depth for angle $i$, $\vec{q}_i$ is the modulation direction, and $\phi_{i,j}$ is the phase offset for angle $i$ and phase step $j$. All SIMFLUX and SIMCODE localizations with $\gamma > 0.1$ were filtered out. In addition, every modulation based estimate that was further away than 0.5 pixels from the fitted 2D Gaussian (in the case of SIMFLUX) or the deep learning position estimate (in case of SIMCODE), was regarded as a non-converging fit, and filtered out as well.

#### 4 DMD pattern generation and design of structured illumination

In the SIMFLUX system, the digital mirror device (DMD) was used as a binary grating (Supplementary Fig. 6). The effective grating pitch size can determine the diffraction angle, which can vary the numerical aperture (NA) of structured illumination. SIMFLUX imaging requires 6 patterns (2 orientations and 3 phase steps) to form a complete imaging cycle. The DMD pattern ($p$) was generated in the following way:

$$r_i = \sin(k_x X + k_y Y + \phi_i) \tag{8}$$

$$p = \begin{cases} 1, r_i \geq 0 \\ 0, r_i < 0 \end{cases} \tag{9}$$

where $k_x$,$k_y$ stands for the pitch wavenumber of the pattern, $X,Y$ stands for the coordinates of a micromirror on the DMD, and $\phi_i$ stands for the phase steps. The wavenumbers $k_x$ and $k_y$ are defined as follows:

$$k_x = 2 \cdot \pi \sin(\theta)/\Delta \\ k_y = 2 \cdot \pi \cos(\theta)/\Delta \tag{10}$$

where $\theta$ is the orientation of the grating pattern on the DMD. SIMFLUX requires two pattern orientations (x and y) which are orthogonal to each other. To generate the x-pattern, we set $\theta$ to be 45°; to generate the y-pattern, we set $\theta$ to be $-45°$. $\Delta$ is the grating period. The DMD chip was rotated by 45° such that the reflected light from the DMD was parallel to the optical table. The choice of $\Delta$ affects the maximum illumination NA. The diffraction angle for the $\pm1$ diffraction orders of the DMD ($\theta_{DMD}$) can be expressed as follows:

$$\theta_1 = \sin^{-1} \frac{\lambda}{\Delta} \tag{11}$$

After the diffracted light passes the first 4f relay (first and second lens) of the SIMFLUX system, the diffraction angle is modified and can be expressed as:

$$\theta_2 = \theta_1 \frac{f_1}{f_2} \tag{12}$$

Then, the third lens focuses the $\pm1$ diffraction orders into the back focal plane of the objective lens. The distance between $\pm1$ diffraction orders at the back focal plane of objective lens ($d$) can be expressed as:

$$d = 2 f_3 \theta_2 \tag{13}$$

The illumination NA ($\mathrm{NA_{ill}}$) can be expressed as follows:

$$\mathrm{NA_{ill}} = \frac{d}{2 f_{\mathrm{obj}}} \tag{14}$$

In the SIMFLUX system presented in this work, a $\Delta$ of 3.977 pixels on the DMD chip was chosen, equivalent to 54.48 $\mu$m. Using the above equations, the illumination NA was found to be 1.47. As the presented system uses a grating pattern with a non-integer number of pixels per pitch, minor diffraction orders around the main diffraction order can be removed using pinholes acting as spatial filters at the plane conjugate to the DMD (i.e., after the first lens, Supplementary Fig. 6). The position of the pinholes corresponding to the $\pm1$ diffraction orders can similarly be calculated from:

$$d = 2 f_1 \theta_1 \tag{15}$$

#### 5 Supplementary figures

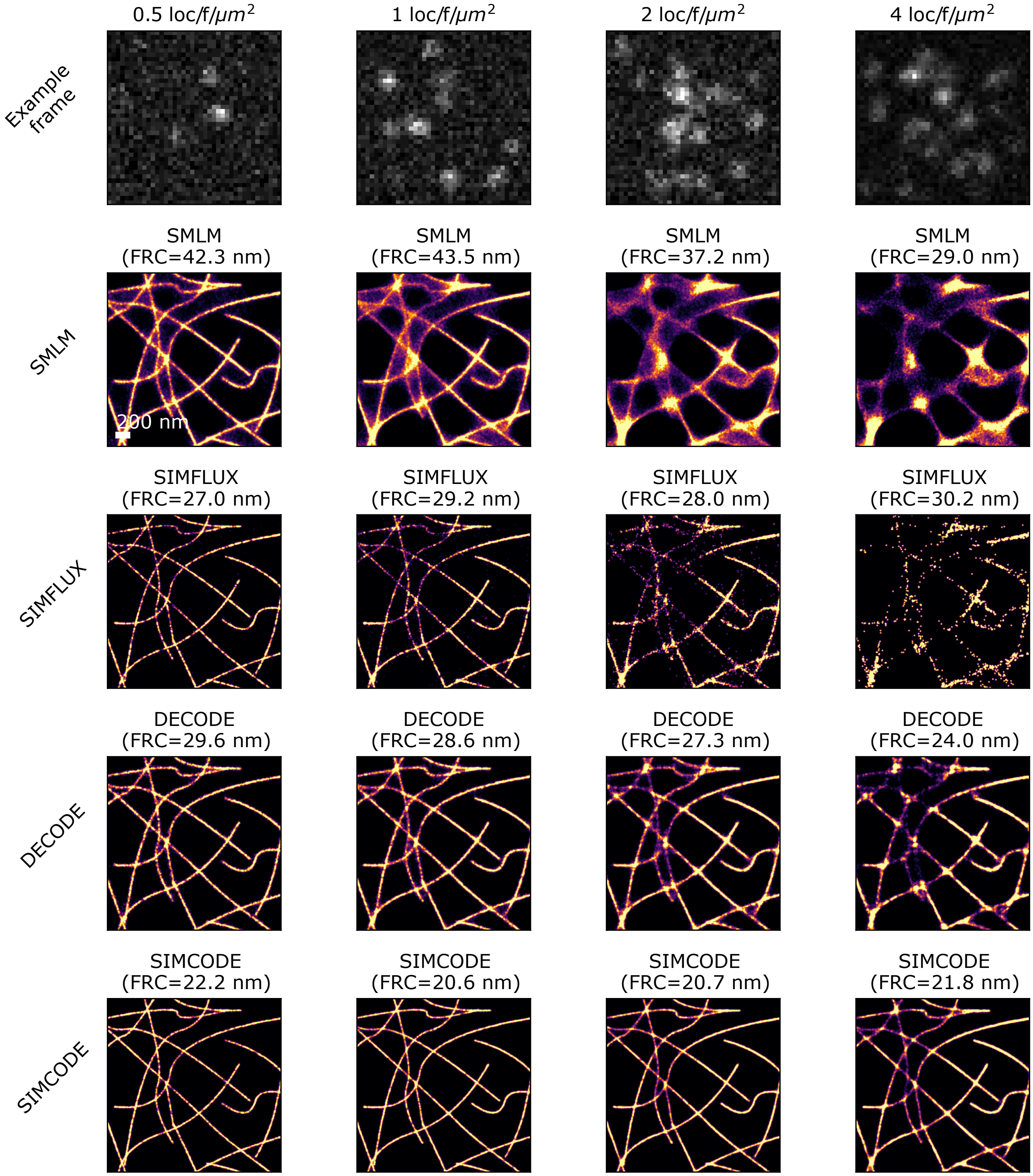


**Figure 1.** Benchmarking the effect of emitter density on reconstruction quality for the different methods. The imaging simulation parameters (such as PSF size, modulation pattern, background, etc.) and ground truth generation was done in the same way as for Fig. 2 in the main text. Emitter density was adjusted by changing the $k_{on}$ and $k_{off}$ values (see Section 3 above) according to the mean on-emitter density. All localizations with a CRLB > 0.2 pixel, or 0.13 nm were filtered out. The Fourier Ring Correlation (FRC) of each reconstruction is displayed, but as can be seen does not always correspond with the visual image quality.

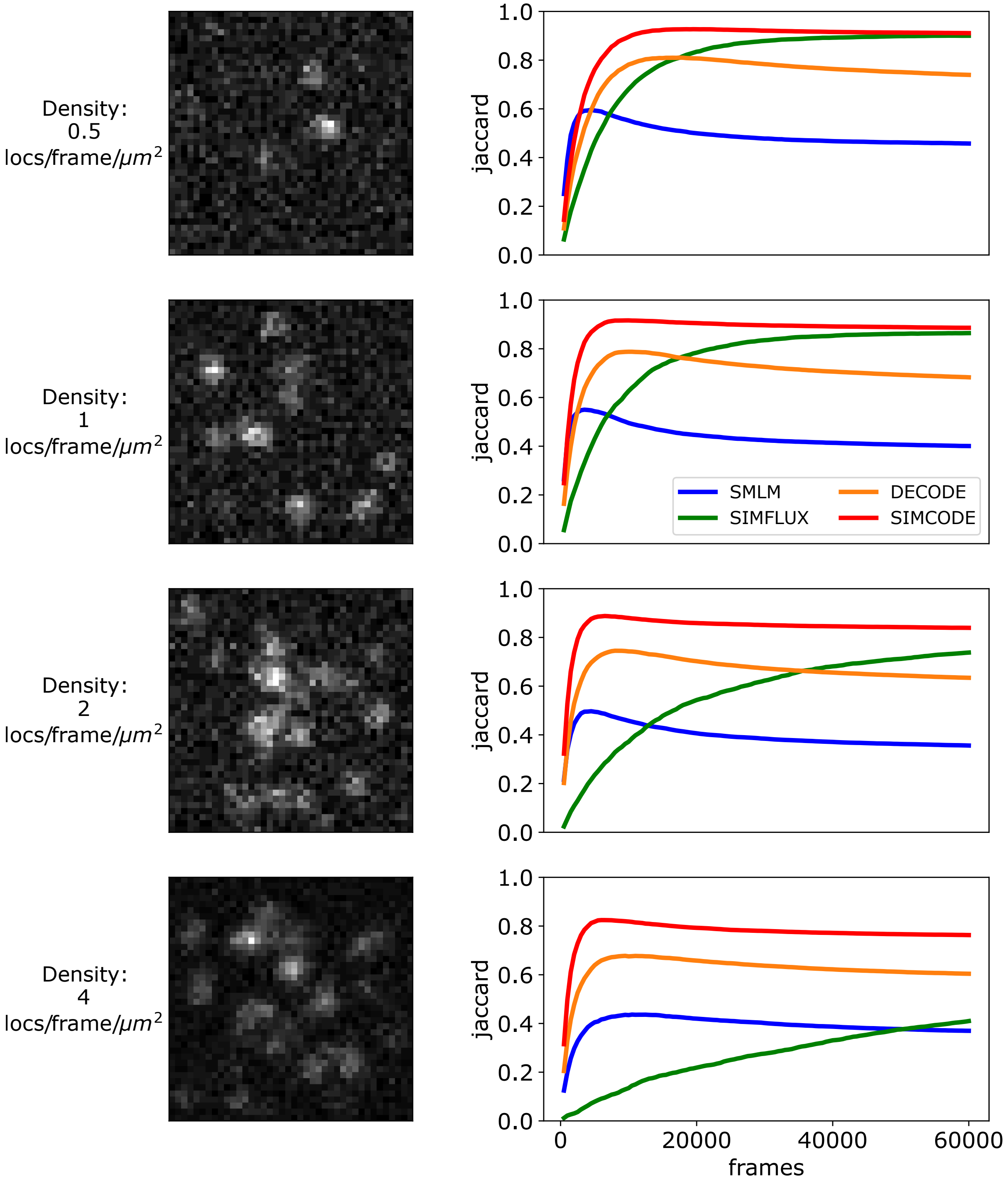


**Figure 2.** To gain insight into the speed of the different localization methods, a Jaccard similarity index was computed between the set of estimated points $A$ and a set of points that make up the ground truth structure $B$. The Jaccard index here is defined as the probability that if a point from the combined set $A+B$ is taken, the point is in both $A$ and $B$ (which can be written as $J(A,B) = p(A)p(B|A) + p(B)p(A|B)$). Inclusion in this point set is defined as any point within 10 nm distance of any point in the set. The Jaccard index is plotted for 4 different densities of active emitters, measuring the similarity between the set of estimated localizations and the set of ground truth points, as localization progresses over time. All settings are the same as for Supplementary Fig. 1.

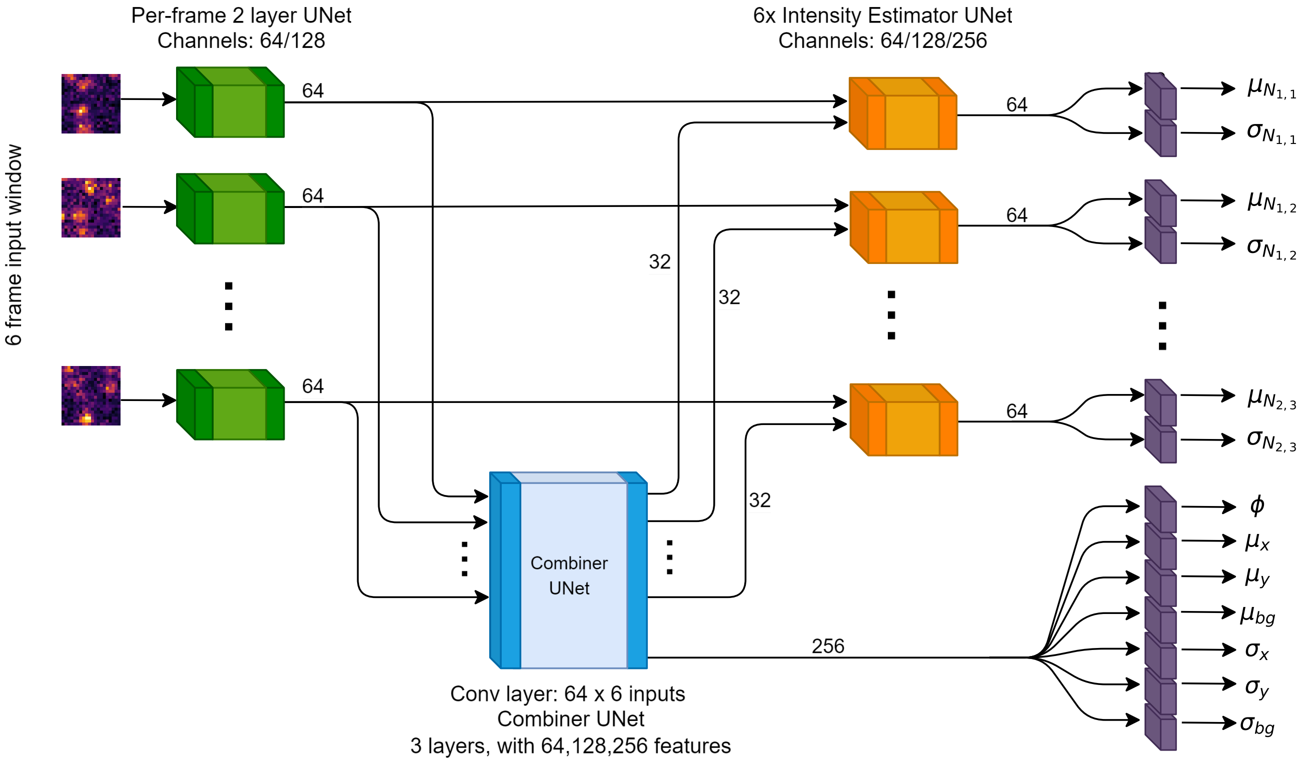


**Figure 3.** Overview of the deep learning model components. The deep learning model is largely a modified version of the DECODE architecture[1], and consists of several U-Net blocks and separate convolutional layers. In the U-Net's, 2D convolutions with stride 2 are used for downsampling, instead of maxpool. Blocks with the same color share weights, except for the purple blocks in the final step. First, the a moving window of 6 raw frames is scaled down to have values between 0 and 1, and are then processed by a U-Net with a depth of 2. The output of this step are 64 values per pixel, which was used as an input for a larger 'combiner' U-Net, that can observe the processed information of 6 frames at a time. The per-frame preprocessing U-Net (green) allows some of the computational burden to be done only once per-frame, instead of repeated for every block of 6 frames, saving some compute. The combiner outputs 256 channels per pixel to the main emitter output module. The combiner U-Net also produced 32 channels per frame (encoding the detected emitters and their positions), which was fed into a U-Net dedicated to estimation of emitter intensities and their associated uncertainties. In total, the network produces an emitter probability $\phi$ as well as mean and uncertainty values ($\mu$, $\sigma$) for the 2D position ($x$, $y$), a background fluorescence at the emitter position ($bg$), and 6 intensities ($N_{i,j}$). Batch normalization was used for all U-Net's. Each U-Net was is also preceded and followed by an extra Conv2D layer, to map between different feature counts. ELU activation functions were used for all non-output activations. The final output blocks (purple) contain a Conv2d layer with kernel size 3, followed by an exponential linear unit (ELU) activation, a linear transformation to go to 1 value, and then either a sigmoid (for emitter probability $\phi$ and parameter $\sigma$ values) or the hyperbolic tangent $tanh(x)$ (for the parameter $\mu$ values) activation. This makes all outputs have a maximum magnitude of one. After this, they are rescaled to physical units by user predefined factors, such as the maximum background or the maximum intensity, depending on the configuration file.

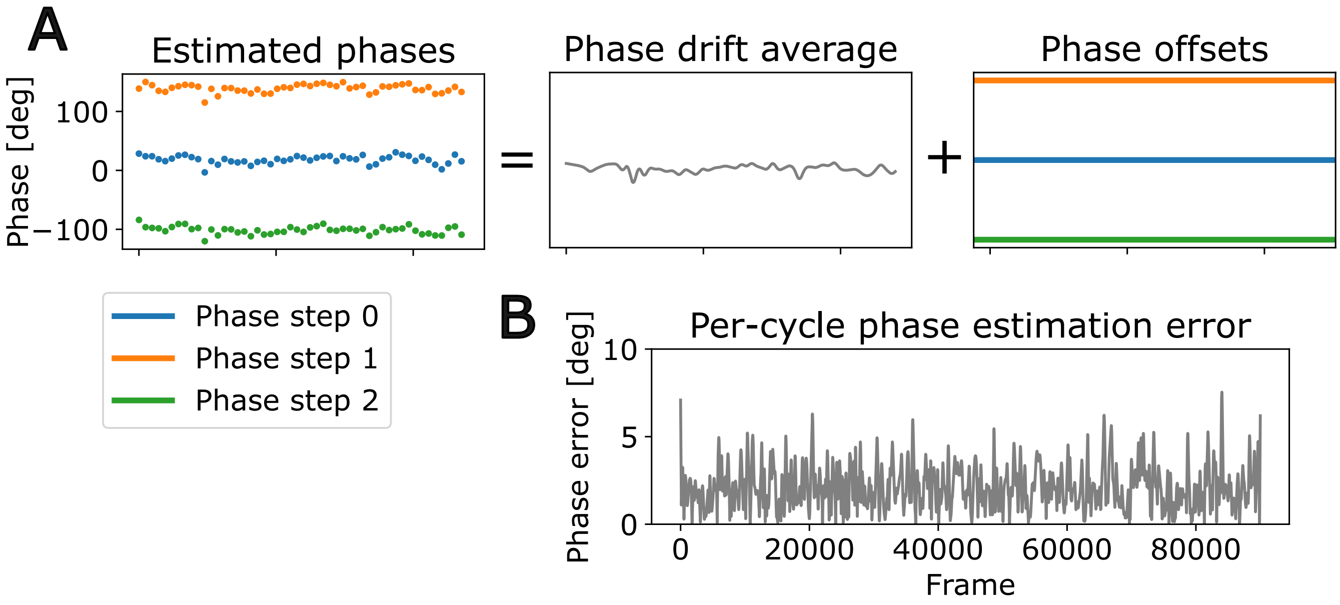


**Figure 4.** For each axis, the phase drift was assumed to be shared over the 3 phase steps (panel **A**). This is a logical assumption considering the use of the DMD with it's discrete pixel steps, and allows us to perform the pattern fits with an improved phase estimate. Panel **B** shows the standard deviation of the phase estimates over time, computed over the 3 phase steps at each imaging cycle. This plot is based on the experimental dataset as shown in main text Fig. 3, processed by SIMCODE. The average phase error was 2.16 degrees, leading to a mean estimated error of $2.16/\sqrt{3} = 1.25$ degrees.

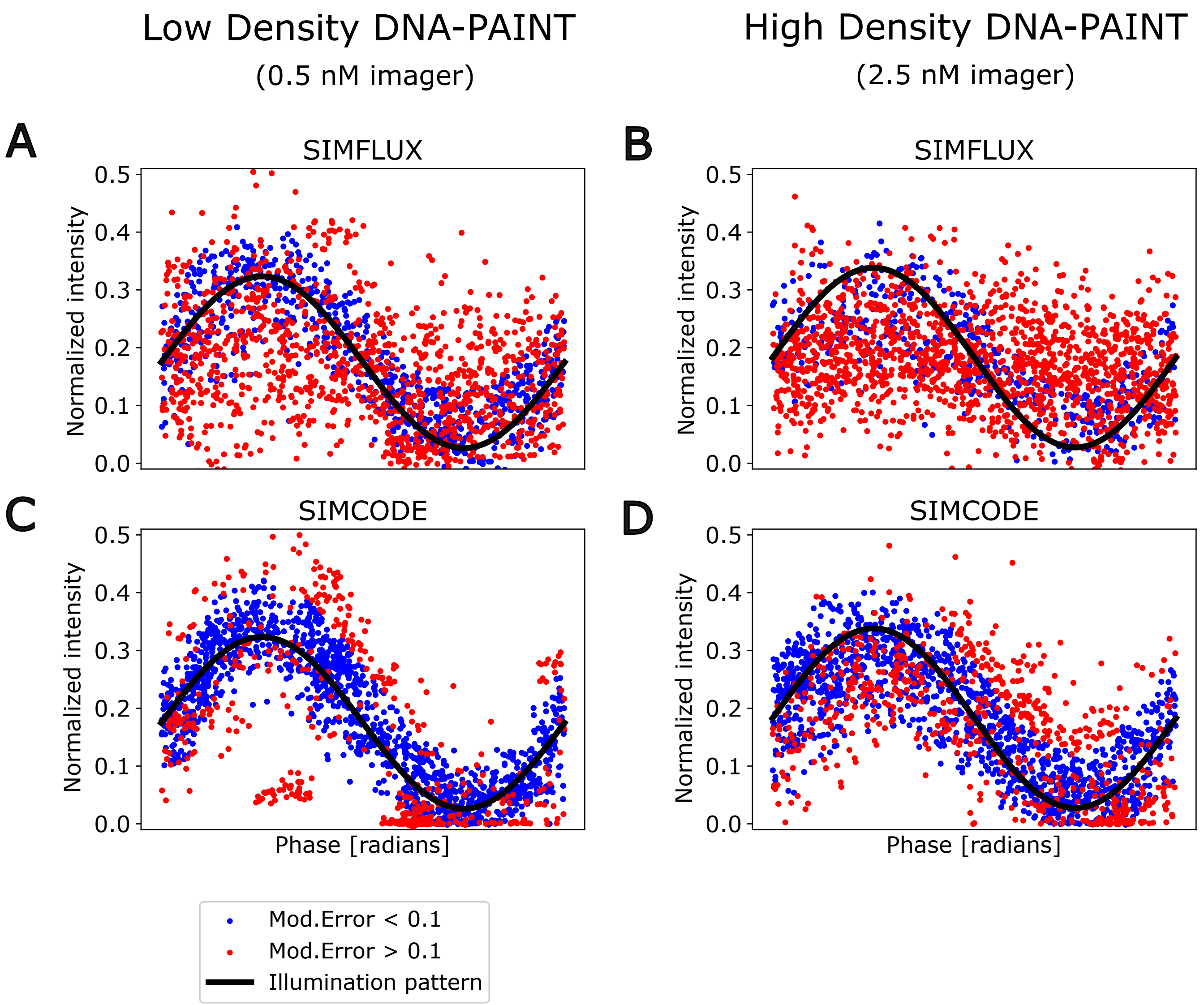


**Figure 5.** These panels show subsets of the localizations and their intensities mapped onto a single period of a modulation sine wave. Intensities were normalized per localization, by dividing by the sum of the 6 intensities $\mu_{N_1} / \sum_i^6 \mu_{N_i}$. The phase as used in the X axes of these plots was computed from the spot's localization-based position estimate (from the SMLM 2D Gaussian fit in panels **A**, **B**, and from the deep learning model in panels **C**, **D**). Depending on the modulation error threshold, localizations were either included in the reconstruction (blue) or excluded (red).

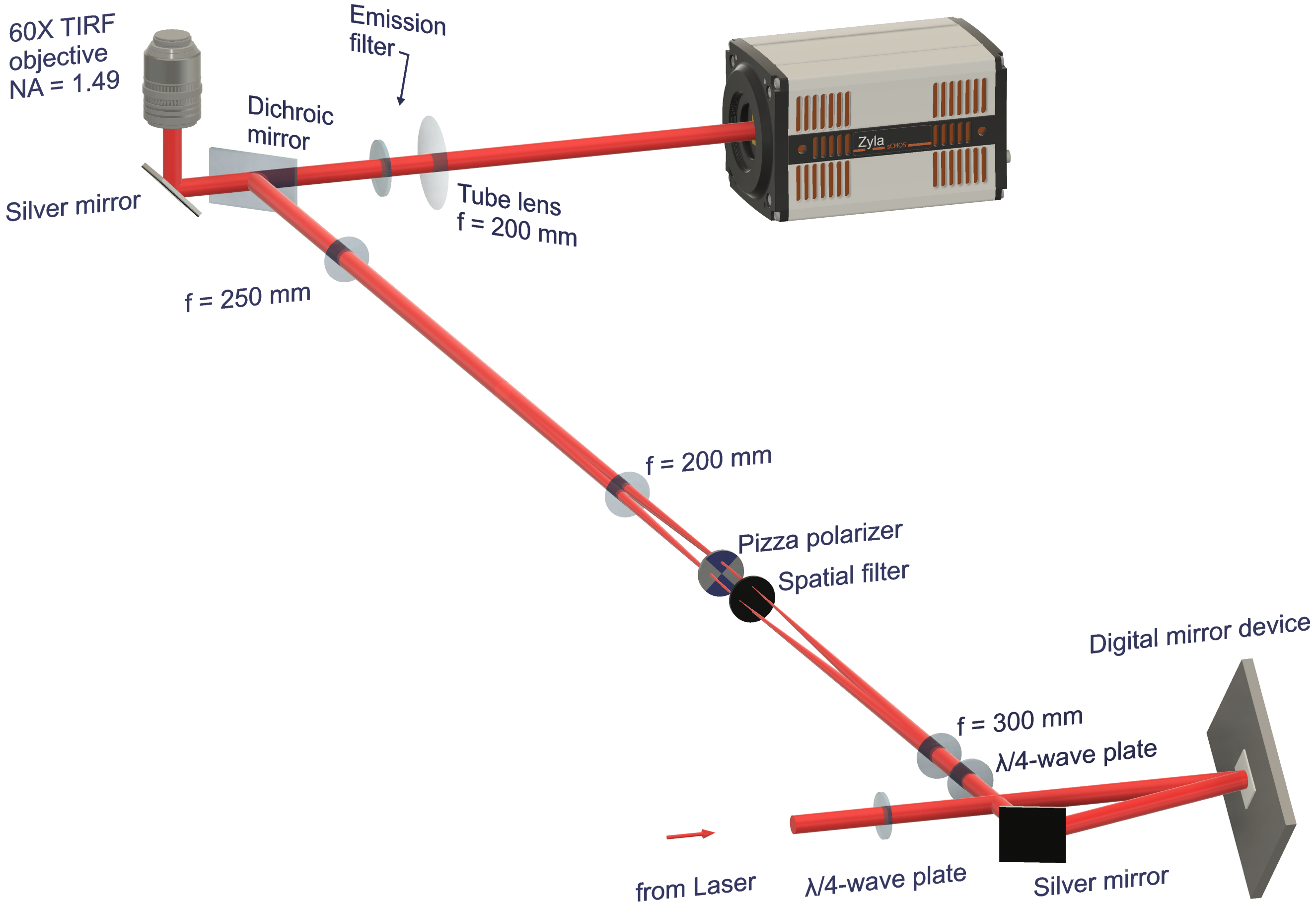


**Figure 6.** 3D rendering of optical setup showing the illumination and emission beam paths. Note the 45 degree rotation of the digital mirror device, the use of $\lambda/4$ plates and pizza polarizers to adjust the modulation contrast of the modulated illumination pattern, and the illumination relay lenses to match the diffracted beams to the numerical aperture of the objective.

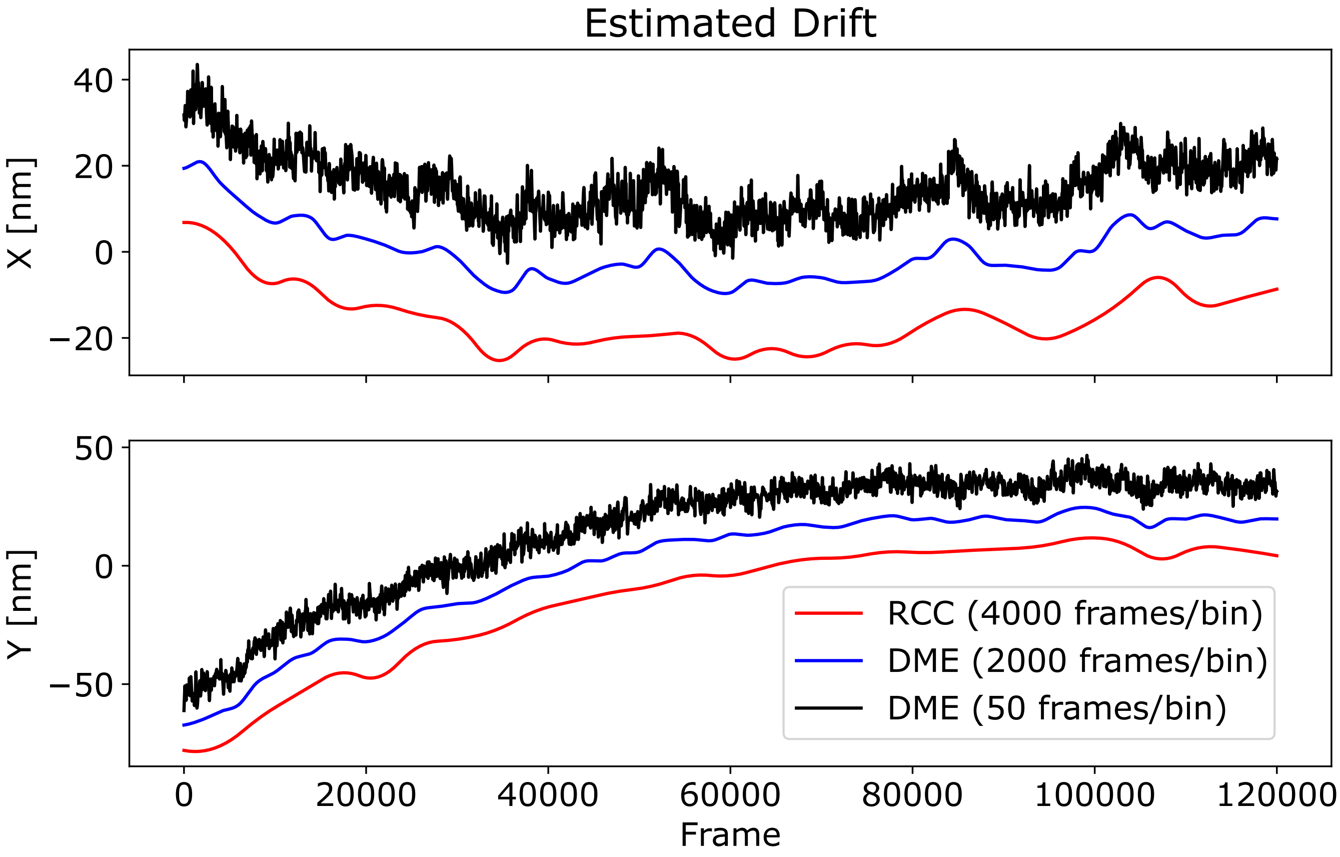


**Figure 7.** Estimated drift for the localizations of main text figure 3 (low density DNA-PAINT). The experimental datasets were processed using Drift at Minimum Entropy (DME)[4] to ensure correction at a high time resolution. The DME implementation only runs well on Windows, and was performed on a RTX 4060 laptop GPU. The drift was estimated on the SIMCODE localizations, and applied to the localizations of the other methods, so the localization method comparison is independent of differences in drift estimation accuracy. To ensure convergence of DME, the localizations were processed in the following way: first, any localizations with a CRLB $> 0.15$ pixels were rejected. Then, redundant cross-correlation (RCC) was performed at a super-resolution of $4\times$ and a binning size of 4000 frames. The top 1M localizations were selected based on CRLB, and DME was run at 2000 frames/bin for a coarse DME estimate. Finally, the coarse estimate was used to run DME at 50 frames/bin (also capped at 1M localizations) to achieve the final drift estimate. Cross validation of the drift estimated using a vertical split of the FOV resulted in a root-mean-squared displacement of 5.0 nm and 4.8 nm for the X and Y axes, respectively, suggesting most of the drift is corrected. However, there could still be some remaining drift or vibration on a short time scale. For visual clarity, the visualization shows the drift traces with a small offset.

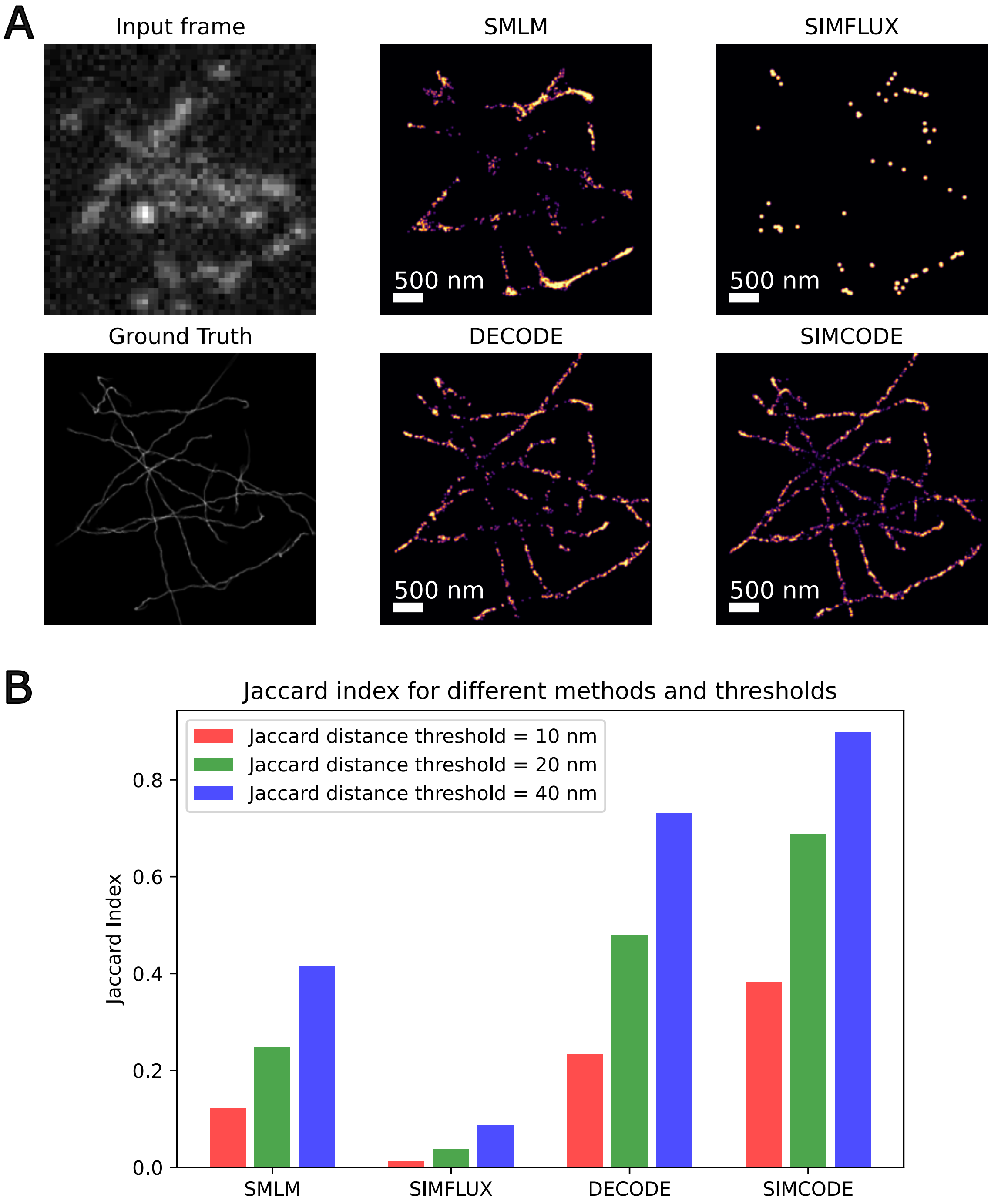


**Figure 8.** We performed reconstructions using the 4 different methods on a simulated moving microtubules. The microtubule simulation was done similar to our density simulations (see Figure 1), but with brownian motion and a spring-like microtubule mechanics added over a series of 60000 frames. The reconstruction was divided into blocks of 1000 frames, with each block of 1000 frames resulting in a single reconstructed frame. One of the input frames, as well as the ground truth for a single reconstructed frame, is shown in panel **A**, as well as the reconstructions for this particular reconstructed frame. A full reconstruction movie can be found on our github repository as well as the supplemental media. To quantify the reconstruction accuracy, we computed the Jaccard index for each method and each reconstructed frame, compared to the ground truth. Panel **B** shows the average Jaccard index for each method, with 3 different distance threshold used for the Jaccard calculation. For the SIMFLUX reconstruction, we used the estimated pattern from SIMCODE, as the points from the SMLM pipeline were too sparse to estimate a modulation pattern.
